# High-Resolution 3D Mapping of Human Cortical Vasculature

**DOI:** 10.64898/2026.05.14.724532

**Authors:** Vanja Ćurčić, Youri Adolfs, Matthias J.P. van Osch, R. Jeroen Pasterkamp, Natalia Petridou

**Affiliations:** Translational Neuroimaging group, Centre for Image Sciences, UMC Utrecht, The Netherlands; Department of Translational Neuroscience, UMC Utrecht Brain Center, The Netherlands; C.J. Gorter MRI Center, Department of Radiology, Leiden University Medical Center, Leiden, The Netherlands; Leiden Institute for Brain and Cognition, Leiden University, Leiden, The Netherlands

## Abstract

A quantitative description of the three-dimensional organization of the human cortical vasculature at micron scale is critical for understanding the influence of the vascular architecture on cerebral blood flow, neurovascular coupling, neurological disease, and human neuroimaging signals. However, high-resolution volumetric vascular data from human cortex are scarce, as the acquisition relies on postmortem microscopy, and tissue-clearing and whole-mount immunolabeling approaches developed for rodent tissues are less effective in heavily crosslinked, pigment rich fixed human cortex. Here we introduce h-iDISCO+, a workflow that enables whole-mount immunolabeling and high-resolution light-sheet imaging of arteries, capillaries, and veins in long-term formalin fixed human brain tissue. By integrating extended oxidative and photobleaching prior to tissue staining, we achieve uniform sample transparency and full-depth antibody penetration. Applying this workflow to tissue samples from human primary visual cortex, we quantified vessel geometrical properties and vascular density across cortical depth. This approach allows quantitative reconstruction of human cortical vascular networks at micron resolution, enabling volumetric datasets of the human cortical vasculature that were previously unavailable.

## 1 INTRODUCTION

The human brain relies on a dense and highly organized cortical vascular network to sustain exceptionally high and continuous metabolic demand, given its limited energy reserves. This network comprises micrometer-scale vessels and exhibits remarkable geometric complexity arising from its three-dimensional (3D) organization. In recent years, increasing attention has focused on how the cortical vascular network supports continuous cerebral blood flow and responds to neuronal activity, as well as how its dysfunction contributes to neurodegenerative disease onset and progression^1–7^.Quantitative descriptions of the 3D cortical vascular organization and topology at microscale are therefore essential for understanding and modeling physiological processes such as cerebral blood flow regulation^5,10^, oxygen transport^8,9^, and neurovascular coupling, as well as for determining the relationship between vascular dysfunction and neurodegeneration^5,11^. Such quantitative knowledge is also increasingly important for the interpretation of human non-invasive functional imaging data, including functional Magnetic Resonance Imaging (fMRI)^12^. Despite its central role in brain health, disease mechanisms, and the neuroimaging signal formation, the 3D organization of the human cortical vasculature is insufficiently characterized. High resolution data are scarce and required postmortem contrast injection approaches, yielding images of cleared thick tissue sections and vascular corrosion casts that support limited quantitative analysis^13,14,15,16^. Consequently, our knowledge on the microscale 3D cortical vascular organization mainly relies on rodent studies, despite distinct differences in vascular anatomy between humans and rodents.

Recent advances in optical clearing, whole-mount immunolabeling, and light-sheet fluorescence microscopy (LSFM) have transformed 3D imaging of the cortical vasculature in mice brain at microscale ^17^. Optical clearing reduces light absorption and scattering, while immunolabeling of vascular-specific proteins enables visualization of arterial, capillary, and venous networks. However, applying these techniques to post-mortem human brain tissue has been challenging^18^. The high degree of myelination, extensive protein crosslinking caused by long-term fixation, and the accumulation of iron-rich heme pigments and autofluorescent lipofuscin in aged human brains impair tissue clearing efficiency and limit antibody penetration. As a result, protocols such as CLARITY^19^, CUBIC^20^, MASH^21^, and iDISCO+^22^, although effective in rodent tissue, often show important trade-offs in human samples between transparency, labeling depth, and imaging quality, highlighting the need for human-specific optimization.

Here we introduce h-iDISCO+, an iDISCO+based workflow that enables vascular immunostaining in long-term formalin-fixed human brain samples. The protocol incorporates an extended oxidative bleaching step combined with photobleaching prior to staining to reduce residual pigmentation without compromising antigenicity, enabling whole-mount labeling of arterial, capillary, and venous compartments. It achieves efficient clearing and vascular immunostaining in 1 cm blocks of post-mortem human brain tissue stored for over a decade, with a total processing time of approximately 30 days. Combined with LSFM and deep-learning-based segmentation, this approach allowed quantitative reconstruction of human cortical vascular networks at micron resolution, enabling volumetric datasets of the human cortical vasculature that were previously unavailable.

## 2 RESULTS

Our h-iDISCO+protocol produced uniform optical transparency in long-term (7-18 years) formalin fixed aged human brain samples, enabling full-depth antibody penetration and LSFM encompassing the entire cortical thickness of the samples (1.1 to 1.8 mm, n=6 samples from the primary visual cortex of two donors with no cerebrovascular pathology (F 68, and M 67 years)). A representative example is shown in Fig. 1A, placed on millimeter-scale paper for size reference. The full workflow for sample preparation and immunostaining including processing times is summarized in Fig. 1A. Formalin fixed samples are initially dehydrated, followed by extended bleaching with 15%hydrogen peroxide (H_2_O_2_) (refreshed daily), combined with photobleaching using LED light at room temperature for 7 days. Samples are then prepared for immunostaining, in which the tissues were labeled with antibodies against Podocalyxin to visualize the entire vascular network and α–smooth muscle actin (αSMA) to identify arteries, which exhibit a higher abundance of smooth muscle cells compared to veins. We first tested the effectiveness of the bleaching approach by evaluating the sample optical transparency after clearing (n = 6 samples, from the primary visual cortex of one donor with no cerebrovascular pathology, F 50 years). We tested different concentrations of H_2_O_2_ (5, 10 and 15 %), at room temperature or 4 degrees Celsius (Supplementary Fig. 1A). Extended oxidative bleaching (7 days instead of overnight used in the original iDISCO+) using 15 %H_2_O_2,_ in combination with continuous application of LED light, gentle shaking and at room temperature led to optimal results in terms of pigmentation reduction and thus the optical transparency achieved after clearing. Due to the low residual pigmentation, light penetration was uniform throughout the depth of the entire sample even while using one-sided illumination (Supplementary Fig. 1 G-H). Next, we tested if the extended bleaching protocol affected the antigenicity of the samples by comparing the h-iDISCO+protocol to iDISCO+(Supplementary Fig. 1 G-L) for 3 tested antibodies (Podocalyxin, αSMA and CD31 (Platelet Endothelial Cell Adhesion Molecule-1 (PECAM-1)). Comparative staining tests confirmed that the extended bleaching with higher concentration of H_2_O_2_ (15%) preserved antigenicity relative to the standard iDISCO+protocol. We performed additional experiments to determine the optimal concentration of the 3 antibodies (Supplementary Fig. 2). CD31 was not used in the final protocol due to poor specificity and noisy signal (Supplementary Fig. 1 K, L, Supplementary Fig. 2 J).

**Figure 1.**
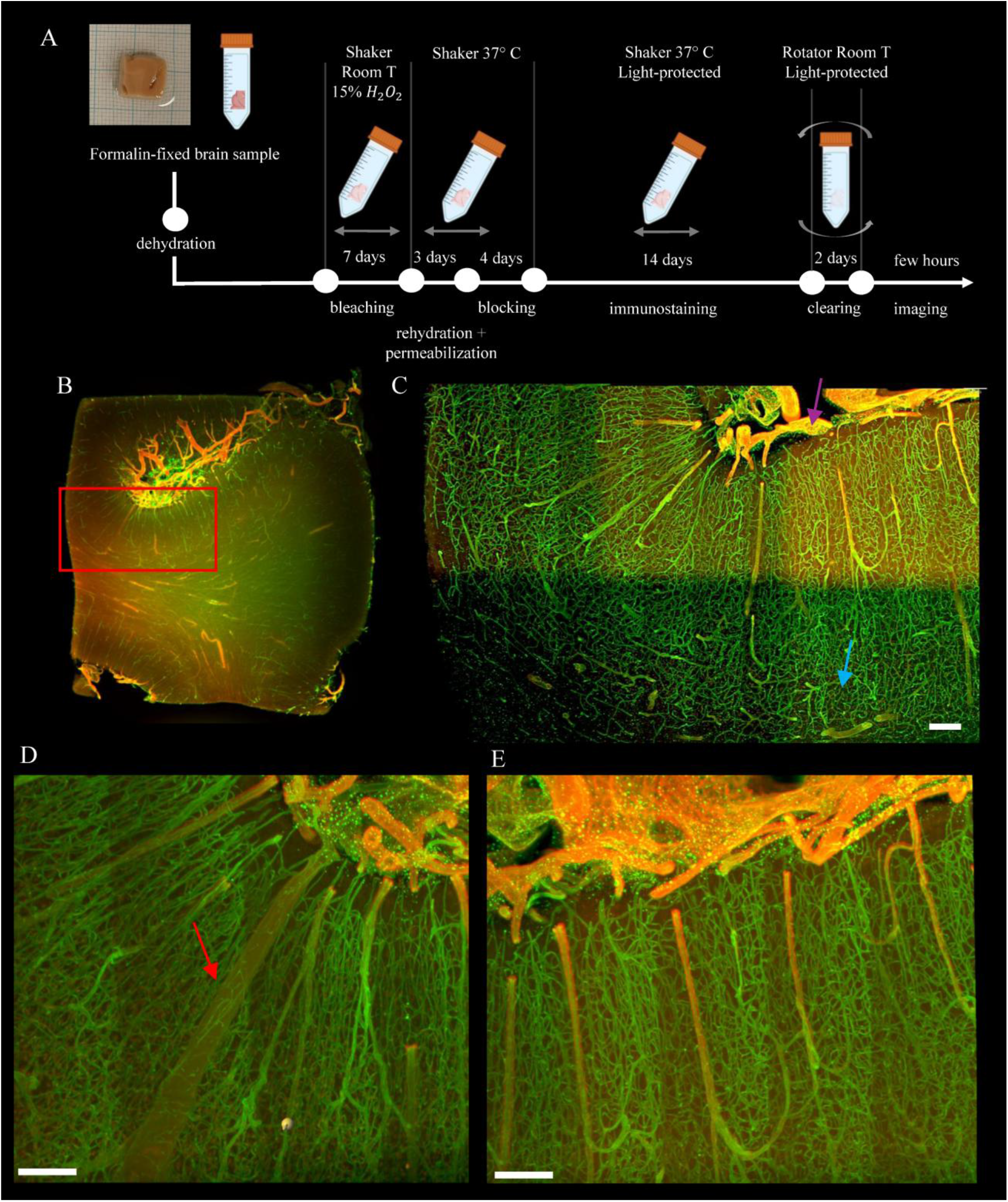
h-iDISCO+protocol for long-term formalin fixed human brain tissue and full vasculature staining results for a representative sample S2. A) Brain sample (S2) used, and protocol steps including their duration. Steps include sample preparation: dehydration, bleaching (15%H_2_O_2_ 7 days at room temperature, shaking under LED light), rehydration and permeabilization, blocking, immunolabeling including incubation with primary and secondary antibodies, clearing, and LSFM imaging. B) Sample overview made for selecting an ROI for high resolution imaging, where the border between GM and WM is visible C) High resolution (0.65 x 0.65 x 2 µm) image of the selected ROI, with purple arrow pointing at the pial surface and blue arrow at the GM/WM boundary. D-E) Zoomed sections corresponding to a 300 µm thick rendering, showing a large intracortical vein denoted with a red arrow (D) and multiple intracortical arteries and dense capillary network (E) Scale bars 200 µm.

After establishing the bleaching protocol and optimal concentrations of antibodies, we mixed Podocalyxin and αSMA to stain the full vasculature. Our results showed that light propagated through the entire sample (1 x 1 x 0.5 cm) (Fig. 1B), and the detected antibodies successfully penetrated the entire cortical thickness (≈2 mm) (Fig. 1C). The applied immunostaining enabled visualization of capillaries and veins (Podocalyxin staining is displayed in green), and arterial compartments (αSMA in displayed in red) (Fig. 1 B-E). Podocalyxin uniformly labeled the entire vascular tree, whereas αSMA staining was strongest in arterial walls, permitting differentiation between penetrating arteries and veins. Low-magnification overviews demonstrated consistent staining across the sample (Fig. 1B). Differences in vascular organization between gray matter (GM) and white matter (WM) can be observed. Arteries and veins in GM are penetrating relatively perpendicularly from the pial surface towards WM, where upon approaching or entering the white matter they change direction and adopt a more tangential orientation, with vessels in WM tending to run parallel to the cortical surface^13,16^. The capillary bed exhibited dense, mesh-like organization.

Regions of interest (ROIs) were selected for high-resolution imaging (red rectangle in Fig. 1B) to capture cortical regions spanning the full cortical depth (Fig. 1C), revealing the complex structure and topology of the vascular tree, including intracortical arteries veins and the dense capillary bed. Vertical band artefacts visible in high resolution images are a consequence of dynamic focusing of the light sheet (see Discussion). To quantitatively assess the vascular morphology, selected ROIs were preprocessed to homogenize illumination differences upon which vessels were segmented using the tUbeNet deep-learning framework^23^. The resulting segmentation predictions showed the vessel geometry, as seen in displayed 100 µm maximum-intensity projections (Fig. 2). These images were further processed to remove noise and fill small holes, followed by binarization. Binary masks were used to compute geometric properties of vessels (diameter, length and tortuosity), (Fig.3) using a custom-built MATLAB pipeline (see Supplementary Material). Vessel geometric properties showed distinct distributions across different vessel classes (Fig. 3). Arterial and venous diameters spanned a broad range (Fig. 3 A,G), while capillaries were identified as vessels with diameter threshold of 10 µm (Fig. 3 D). Segment lengths were strongly right skewed across all classes (Fig. 3 B, E, H), with the longest tails observed for arteries. Tortuosity values were generally close to 1.2, indicating low curvature, with sample-to-sample variability most evident in the microvascular compartments (Fig. 3 C, F, I). Next, we quantified vascular density, defined as the volume fraction of segmented vessel voxels, as a function of cortical depth in successive 25 µm depth bins (Fig. 4). Profiles were extracted in the direction from the pial surface (PS) to the GM-WM boundary. Cortical depth was normalized to values between 0 (PS) and 1 (WM). Red dashed boxes indicate the regions in which density was computed, selected so that so that cortical curvature was homogenous within each region. Graphs (Fig. 4) show initial high values corresponding to high vascular density at the pial surface, a dip corresponding to the space between pial vessels and cortex, followed by an increase in density through cortical depth, with highest values occurring at depths from 0.5 to 1mm depth depending on the sample.

**Figure 2.**
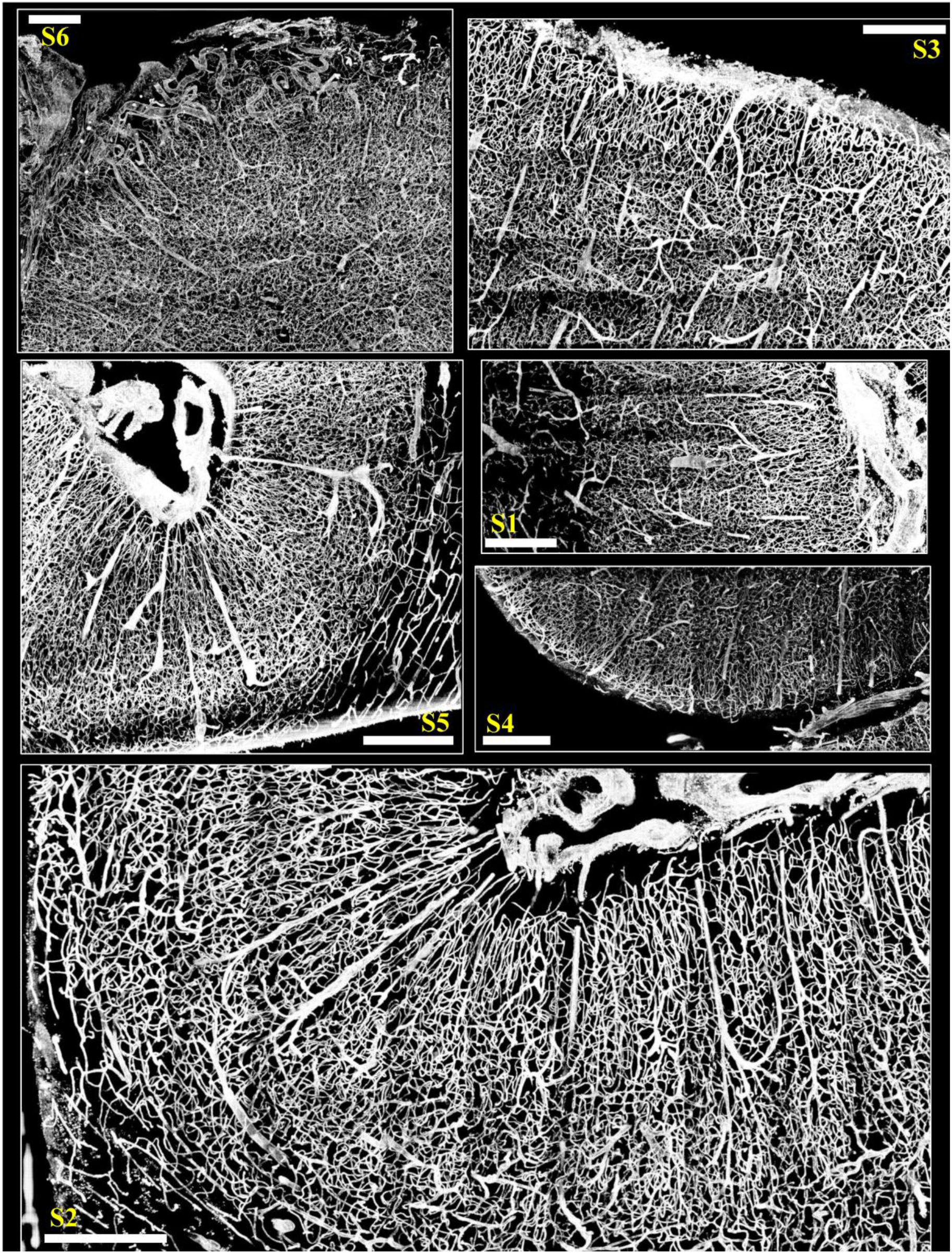
tUbeNet vessel probability maps across samples (S1–S6). The panels show representative 100 µm maximum-intensity projections of vessel probability maps predicted by tUbeNet, in which each voxel is assigned a probability of belonging to the vasculature. The probability maps were subsequently post-processed and binarized for quantitative analysis. Scale bars: 500 µm.

**Figure 3.**
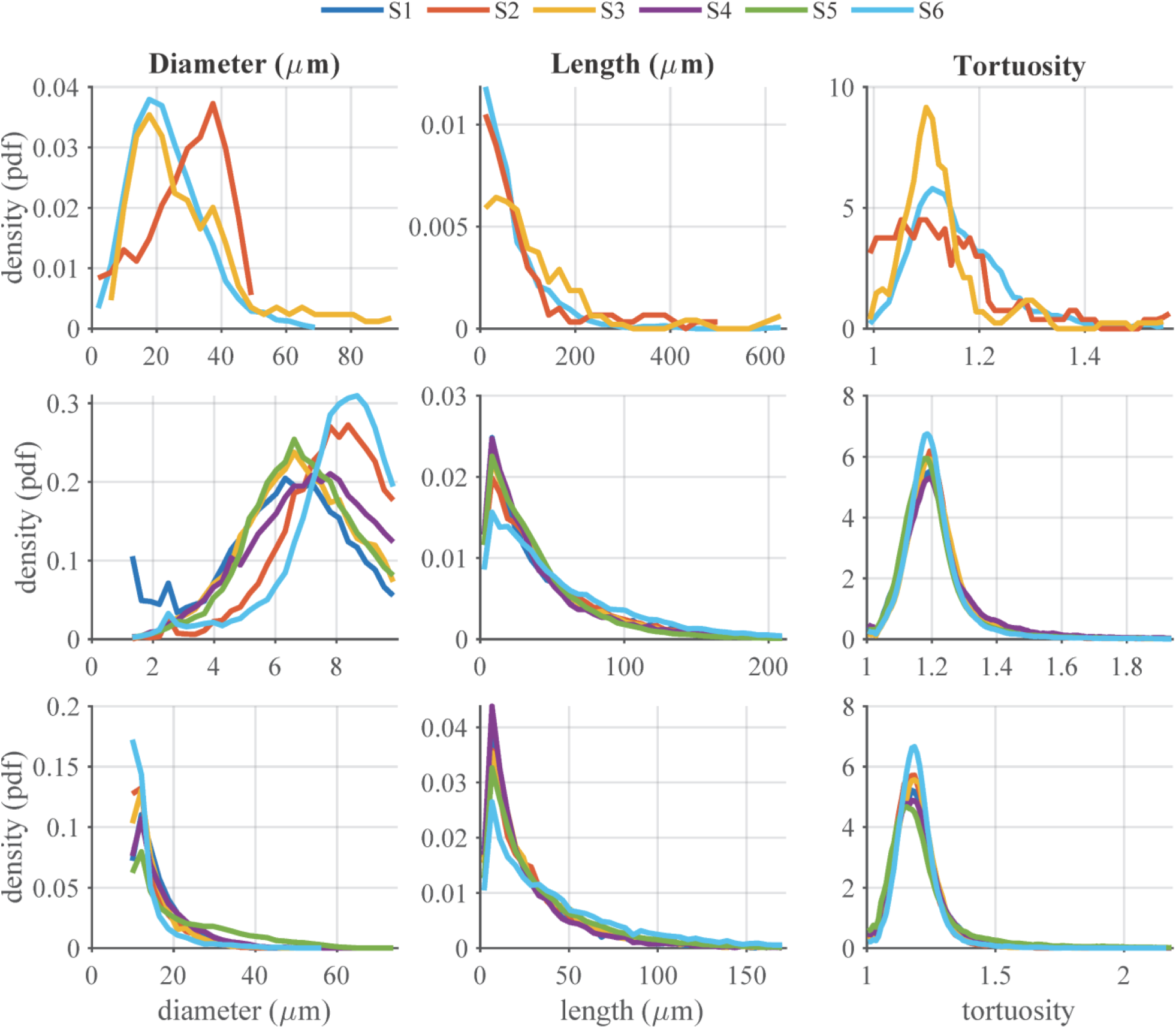
Distributions of vessel geometric properties across vessel classes and samples. Probability density functions of diameter, segment length, and tortuosity for arteries (A–C, top row), capillaries (D–F, middle row), and veins (G–I, bottom row) across six samples (S1–S6, different colors). Tortuosity was calculated as the ratio of centerline path length to straight end-to-end Euclidean distance. All distributions are shown as normalized densities.

**Figure 4.**
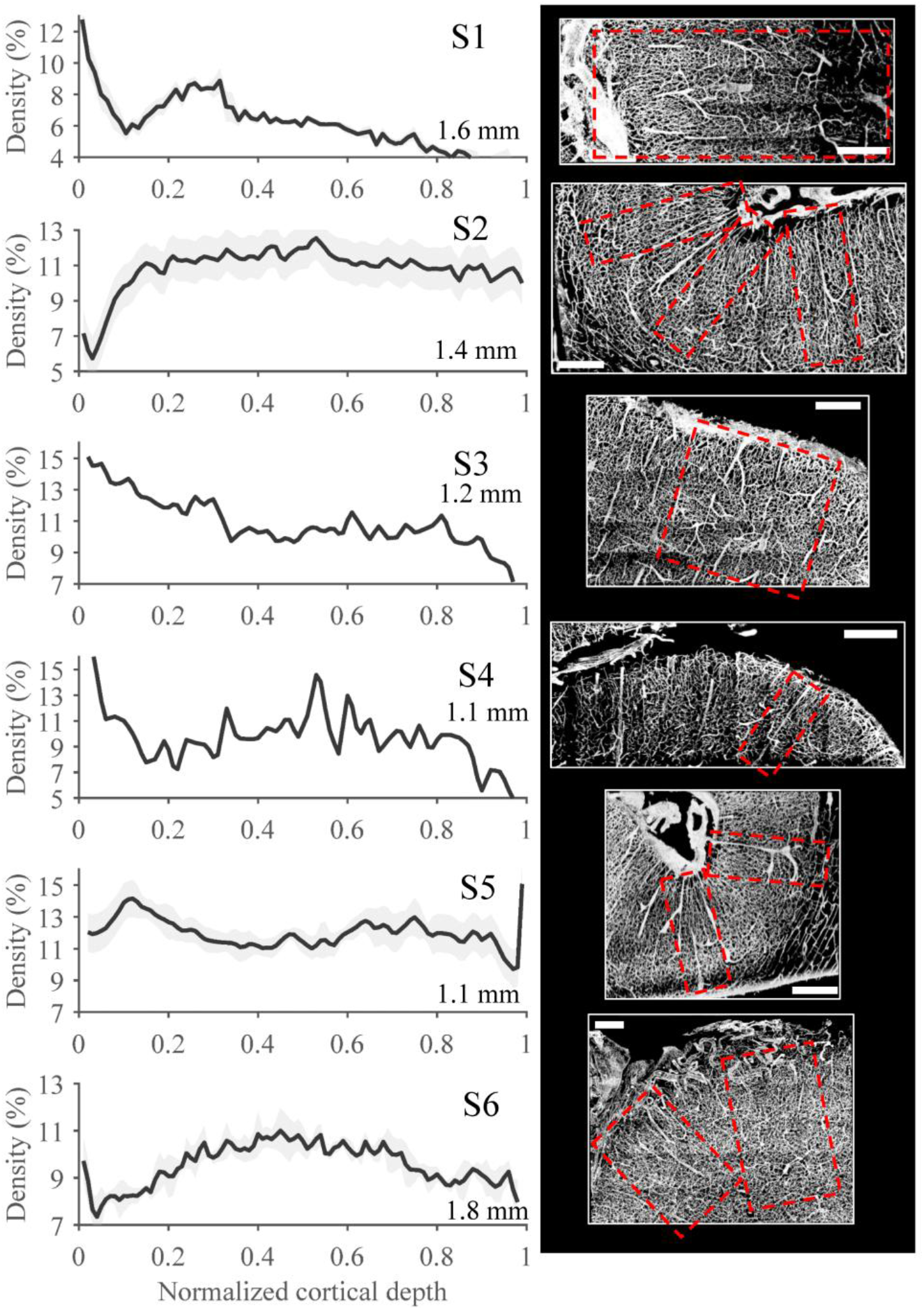
Depth-resolved vascular density profiles across samples. Left: vascular density (vascular volume fraction, %) plotted as a function of normalized cortical depth from the pial surface (PS) to the white matter boundary (WM) for six samples (S1–S6), with denoted cortical depth in mm. Density was calculated in consecutive 25 µm depth bins as the fraction of segmented vascular voxels within each bin. Right: Representative 100 µm maximum-intensity projections for each sample. Red dashed boxes indicate the position of the 3D regions from which density profiles were computed, chosen to ensure relatively homogeneous cortical curvature within each region. For samples with multiple analyzed regions, density values were normalized and averaged, where the shaded band indicates the standard deviation across regions at each depth. Cortical depth was normalized between 0 (PS) and 1 (WM) for all samples.

## DISSCUSSION

We introduce h-iDISCO+, a workflow for clearing, immunolabeling, and high-resolution 3D imaging of the human cortical vasculature in long-term formalin-fixed aged brain tissue. By combining extended oxidative bleaching with photobleaching, and vascular-specific immunolabeling, h-iDISCO+enabled deep and uniform labeling of arteries, capillaries, and veins across the full cortical depth in cleared tissue (Fig. 1). This enabled volumetric reconstruction of the human cortical vascular network at micron resolution, extending capabilities previously limited to mice, and overcoming the limitations of earlier methods applied in human tissues. Importantly, the workflow is compatible with postmortem human brain tissue samples stored for more than a decade. This substantially expands the range of tissues accessible for 3D vascular analysis by eliminating the need for vascular injection-based labeling shortly after death. Using this workflow, we generated volumetric datasets of the human cortical vascular network in the primary visual cortex and quantified geometric properties of arteries, capillaries, and veins. These results advance our quantitative knowledge of the human 3D vascular organization and establish a foundation for interpretation and modeling of the link between the human cortical vasculature and cerebral physiology.

The clearing, immunolabeling, and imaging workflow presented is applicable to long-term stored brain samples, which enables future studies across a wide range of anatomical regions, storage conditions, and disease states, by leveraging the large donor populations available in brain banks. For the present study, we included both male and female donors without cerebrovascular pathology with short post-mortem delay (PMD, 3.5 to 5 hours). Post-mortem delay (PMD) proved to be a major predictor of vascular labeling quality and overall protocol success. In pilot testing, samples with longer PMDs (10.5 hours) resulted in unsuccessful staining, suggesting that a short PMD preserves vascular integrity and enables reliable immunolabeling.

A major obstacle in post-mortem human brain clearing arises from residual pigmentation and extensive protein crosslinking resulting from prolonged fixation. We addressed this challenge with a combination of extended bleaching duration of 7 days, increased H_2_O_2_ concentration of 15%at room temperature, daily refreshment of the H_2_O_2_, and photobleaching, which effectively reduced pigmentation thereby improving optical homogeneity throughout the tissue. The enhanced transparency allowed uniform light penetration even with one-sided illumination and markedly improved image quality compared to the iDISCO+protocol established in mice (Supplementary Fig. 1). Comparative staining demonstrated that the more extended bleaching did not compromise antigenicity relative to the standard iDISCO+protocol (Supplementary Fig. 1). iDISCO+bleaching (overnight 5%H_2_O_2_ in methanol at 4 degrees Celsius) proved insufficient to eliminate pigments in our samples, resulting in uneven transparency, limited light penetration and light scattering (Supplementary Fig. 1). The extended bleaching treatment therefore provides a practical solution for removing residual pigments in aged human tissue while preserving the ability to perform staining with fluorescent antibodies.

Staining with selected vascular specific markers enabled visualization of the entire vascular network (arteries, capillaries and veins) while covering the complete cortical depth (Fig. 1 B-E). In our double-staining approach, Podocalyxin provided complete coverage of the vascular tree, while αSMA stained larger intracortical arteries, although αSMA-labeling was not consistently sufficient to provide automatic discrimination between arteries and veins, as was reported previously in mice^17^. Based on the acquired volumetric data, we quantified vessel diameter, segment length, tortuosity, and vascular density through the cortical depth for penetrating arteries, capillaries and penetrating veins. These metrics were generally consistent with previous reports from perfused or partially reconstructed human datasets^13,15,16^, supporting the validity of our approach. Computed diameters of intracortical arteries fell within the range of ∼50 µm (Fig. 3A). The intracortical venous diameter distribution reached ∼80 µm (Fig. 3G), which falls within the reported range for smaller intracortical veins (∼20 - 65 µm range^13^), although below the upper values reported for the largest intracortical draining veins (up to ∼120 - 125 µm). This might be explained by fixation-induced deformations of veins due to the thin structure of their walls, and segmentation-related splitting of large lumen veins (such as the large intracortical vein in Fig. 1D) into two thinner vessels. Within the defined capillary compartment (diameters <10 µm), the diameter distribution peaked in the range of ∼5.9 - 6.6 µm (Fig. 3D), and the mean capillary segment length ranged between 40 and 58 µm (mean ∼53 - 63 µm in earlier reports, depending on region^15,16^). Segment length estimates are sensitive to the underlying skeleton topology and to how vascular segments are defined. We defined segments as arc length along the centerline between nodes that represent either bifurcations or endpoints, consistent with prior studies based on India ink-perfused datasets. In addition, we provide a quantitative assessment of vascular tortuosity, on the order of 1.2 (Fig. 3C, F, I), a metric of particular relevance given its association to pathological processes linked to Alzheimer’s disease (AD) and Cerebral Amyloid Angiopathy (CAA)^5,11,24^. Furthermore, the computed vascular density exhibited layer dependent values ranging from ∼5%to ∼15%across the cortical depth (Fig. 4). These values are higher than those reported in earlier confocal reconstructions of 300 µm sections of India ink–perfused tissue ^16^ , but in line with a recent 3D large-volume microscopy report in primary visual cortex ^25^. As vascular density is region-specific ^13^, such differences may arise when comparing across studies. Alternatively, these discrepancies may arise from methodological differences in how the vascular density was defined and calculated. In volumetric light-sheet based analysis, density was defined as the fraction of voxels labeled as vessels during segmentation, whereas confocal-based reconstructions estimated lumen volume fraction which might underestimate vascular density due to incomplete perfusion of the capillary network, and limited physical volume sampled by acquisition. Overall, our results provide volumetric, quantitative characterization of the human cortical vascular organization and topology across cortical depth, advancing current descriptions.

Several limitations should be considered. First, pial vessels were incompletely stained, due to antibodies binding to the vessel wall, which lead us to exclude the pial surface from quantitative analysis of vascular properties. In the αSMA channel, skeletonized pial surface vessels formed one connected component, which was excluded prior to the vessel analysis. In the Podocalyxin channel, only regions below the pial surface were selected for analysis by cropping the dataset accordingly. Second, staining of large lumen veins often resulted in hollow-tube appearance which may lead to artificial splitting of a single large vessel into two parallel structures, affecting diameter distributions for large vessels. For this reason, we could not reliably segment arteries in the αSMA channel for samples S1, S4 and S5 (Fig. 3 A-C), which were therefore excluded from the analysis. Third, large vein diameters may have been underestimated because venous structures are susceptible to deformation during fixation, and solvent-based tissue clearing methods such as iDISCO+can additionally induce moderate tissue shrinkage through methanol dehydration and delipidation steps^18^. Fourth, αSMA-labeling was not consistently sufficient for automatic discrimination between arteries and veins^17^. To further enhance labeling quality, future work could test brighter, more photostable fluorophores (e.g., Alexa Fluor Plus conjugates) to improve signal. In addition, directly conjugated primary antibodies may reduce reliance on secondary antibodies, with the trade-off of reduced amplification compared with secondary-based detection. Prolonging incubation times could also improve labeling uniformity but would further extend the already lengthy staining process.

From a technical perspective, large-scale LSFM of human samples requires a balance between field of view, spatial resolution, acquisition time and data size. Because the axial (z) resolution of the light sheet is limited by its thickness, samples were oriented so that cortical depth lay within the higher resolution xy plane, improving image sharpness across cortical layers while reducing the total imaging volume. This setup also ensures that the depth which the emitted light must propagate through the sample to reach the objective is kept as short as possible, thereby reducing scattering, refraction and absorption of emission light^26^.

Additionally, signal intensity in long-term fixed tissue is inherently limited by low antibody antigen affinity. To achieve sufficient signal, higher laser power or longer exposure times are often required, which can accelerate photobleaching. The resulting loss of fluorescence explains the signal decay observed between imaging tiles on the top and bottom in Fig. 1 C. Stripe-like artifacts in the xy plane are caused by the UltraMicroscope Blaze (Miltenyi Biotec) linear blending algorithm. When the thinnest part of the light sheet is dynamically focused several times over the field of view to achieve optimal xy resolution, right and left image are linearly blended in each dynamic focusing point. This leads to decrease of signal standard deviation in the blending range, which will appear as a darker band on images. The artefact becomes more pronounced as z is increasing and when high exposure times are used. This effect was not caused by misalignment and correction of the horizontal focusing, as the microscope was calibrated using the alignment tool and chromatic offset of the horizontal focus was corrected prior to acquisition. Stripe artefacts were mitigated by image preprocessing. Fast tiling with decreased numerical aperture (NA) provided an alternative acquisition strategy that eliminated the need for dynamic focusing and reduced stripe-like artifacts (Fig. 2, S5, and S6). This strategy increased imaging speed and reduced laser energy deposition due to higher sheet thickness, thereby decreasing photobleaching and improving image quality, at the cost of increase of voxel size in the z direction (from 2 to 3 µm).

In summary, we present a workflow that enables high-resolution, full cortical depth 3D reconstruction of the cortical vascular network in long-term formalin-fixed human brain tissue. The workflow extends volumetric vascular imaging from rodent models to the human brain by overcoming key barriers, such as insufficient antibody penetration, residual pigmentation, and incompatibility with aged human tissue, without requiring post-mortem perfusion and without compromising antigenicity. By combining vascular-specific immunolabeling with light sheet fluorescence microscopy, we generated micron-resolution volumetric datasets of the human cortical vascular structure and topology. These datasets reveal the human cortical vasculature in its complexity and provide a foundation for linking the cortical vascular organization to cerebral physiology, disease mechanisms, and neuroimaging signals.

## 4 METHODS

Post-mortem human brain samples were obtained from the Netherlands Brain Bank. Samples were retrieved from long-term formalin-immersed sliced brain samples and further cut to include the primary visual cortex. Neurologically healthy donors were chosen to minimize vascular comorbidities. We used samples from three donors (F 50, F 68, M 67, F 65 years), with post-mortem delays of 5.5, 3.5 4.5 and 10.5 hours. Samples from one donor (F 50) were used for developing the bleaching and clearing protocol. Samples from two donors (F 68 and M 67) were used for immunostaining and analysis of vascular properties and density. Sample from donor F 65 was excluded from analysis due to long PMD (See Section 4.1.2).

### 4.1 iDISCO+protocol for immunostaining blood vessels in human brain samples

#### 4.1.1 Tissue preparation

Each sample was initially dehydrated with Methanol (MeOH), in 6 steps with a gradual increase of MeOH percentage (from 20%, 40%, 60%and 80%) and finishing with two washes in 100%MeOH, where each step lasted for 1 hour and was done in a rotator at room temperature at 20 rpm. Next, the sample was immersed in 33%MeOH and 66%dichloromethane (DCM, Sigma Aldrich cat: 5895810250) overnight in a rotator at room temperature set at 20 rpm, for delipidation. Subsequently, the sample was washed 2 times for 1 hour in 100%MeOH.

These steps were followed by bleaching to remove tissue pigmentation. We evaluated the efficacy of tissue clearing 1) In the standard iDISCO+bleaching step, samples were first chilled on ice for 30 minutes and then immersed in pre-cooled 5%H_2_O_2_ diluted in MeOH overnight at 4 degrees Celsius. After standard iDISCO+clearing. 2) To address residual pigmentation, we implemented a more aggressive bleaching protocol. Samples were bleached for 7 days in 15%H_2_O_2_, with the solution renewed every 24 hours, on a shaking platform at 40rpm under LED illumination (Manutan, Cat A23091, 5 V, 10 W, 1000 lm) at room temperature. The use of LED light was adapted from protocol for clearing heavily pigmented human eyes^27^, and histopathological protocols for highly pigmented melanoma samples^28^.

After 7 days of bleaching, the sample was rehydrated with gradually decreased MeOH percentage (starting from 80%, 60%, 40%, and 20%MeOH). Subsequently, the sample was immersed in Phosphate Buffered Saline (PBS) for 1 hour and then washed 2 times for 1 hour in PTx2 (PBS +Triton X-100 (0.2%) and 0,01%thimerosal). This was followed by a permeabilization step that lasted for 2 days in a solution of PTx.2, 20%DMSO, 2,3%Glycine 0,01%thimerosal, at 37 degrees Celsius in a shaking incubator at 70 rpm. After the permeabilization step, the sample was immersed in the blocking solution (composed of PTx.2, 6%Donkey Serum, 10%DMSO and 0,01%thimerosal) for 4 days at 37 degrees Celsius in a shaking incubator at 70 rpm. After 4 days of blocking incubation, immunostaining was performed.

#### 4.1.2 Immunostaining

We initially selected 3 primary antibodies targeting proteins expressed by endothelial and smooth muscle cells in blood vessel walls (proteins Podocalyxin, Alpha Smooth Actin (αSMA), CD31). Antibody to Podocalyxin was used to stain the entire vascular network. Antibody to αSMA was expected to be more expressed in arteries compared to veins, due to the higher presence of smooth muscle cells in arteries and could ultimately be used to differentiate between arteries and veins. CD31 was expected to provide deeper labeling of endothelial cells in combination with Podocalyxin. We optimized antibody concentrations for each vascular protein marker by testing different dilutions in pilot experiments. These experiments further showed that immunostaining was unsuccessful in tissue sample with a postmortem delay (PMD) of more than 10 hours (F 65). For Podocalyxin (Bio-Techne, AF1658), three concentrations were evaluated: 1:500 (low), 1:200 (medium), and 1:40 (high). For αSMA (Abcam, ab5694), two concentrations were tested: 1:1000 (low) and 1:200 (high). For CD31 (Bio-Techne, AF3628-SP), two concentrations were tested: 1:300 (low) and 1:65 (high). Each antibody was tested independently to determine the concentration for optimal staining. Antibodies against Podocalyxin and CD31 were detected using donkey anti-goat IgG Alexa Fluor 647 (Thermo Fisher Scientific), while αSMA was detected using donkey anti-rabbit IgG Alexa Fluor 568 (Thermo Fisher Scientific), each diluted 1:750. Following individual testing of all three primary antibodies, we excluded CD31 due to strong nonspecific background, which resulted in a noisy signal. Since CD31 and Podocalyxin would be stained and visualized in the same channel, the inclusion of CD31 risked compromising the Podocalyxin signal. We therefore optimized the staining by using both Podocalyxin and αSMA at a final dilution of 1:200 and combined these two antibodies for dual labeling in the final protocol.

#### 4.1.3 Imaging parameters and imaging

Samples were imaged with an Ultramicroscope Blaze (Miltenyi Biotec) light-sheet microscope equipped with Neo sCMOS camera (2048 ×2048 pixels, 6.5 x 6.5 µm pixel size) and controlled with Imspector software (v7.8.1). Imaging was performed using a 4 x numeric aperture (NA) 0.35 MI PLAN objective lens with a dipping cap for organic solvents and 2.5 x post magnifier zoom, resulting in a total magnification of 10x and an effective pixel size of 0.65 µm in specimen plane and a maximum resolution of 1.3 µm at detector. The light sheet was generated with a sheet NA of 0.186 (corresponding to an estimated sheet thickness of 3.9 µm), and image stacks were acquired with a z-step of 2 µm, resulting in a voxel size of 0.65 ×0.65 ×2 µm^3^. Horizontal dynamic focusing of light-sheet was used with optimized number of steps. As an alternative acquisition strategy, fast tiling with reduced sheet NA of 0.061 (corresponding to an estimated sheet thickness of 6µm) was used to eliminate dynamic focusing and thus reduce stripe-like artifacts (Fig. 2, S5, and S6). This approach increased imaging speed and, by generating a thicker light sheet, reduced laser energy deposition, thereby decreasing imaging-induced photobleaching, and improving image quality, at the expense of a larger voxel size in the z-direction (resulting voxel size of 0.65 ×0.65 ×3 µm^3^). To minimize shadowing and maximize signal in the target cortical region while decreasing scanning time, single-sided illumination was used, selecting the illumination side closest to the region of interest, i.e. the cortex. A white light superK Fianium laser FIU-15 (NKT Photonics) at 62%power was used in combinations with the following filters: 1) Ex560/40, EM 620/60 (for Alexa 568) with the laser attenuator at 80%and 500 ms exposure time 2) Ex630/30, EM 680/30 (for Alexa 647) with the laser attenuator at 80%and 1000 ms exposure time.

#### 4.1.4 Data processing and analysis

Raw data from the microscope was converted to Imaris format (version 10.1, Oxford Instruments, RRID:SCR_007370) for visualization and stitching (Imaris Sticher 10.1). Size of selected ROIs for high resolution imaging are given in Table 1. To correct for illumination inhomogeneities across the imaged volume, we applied a custom Python-based flat-field correction in which a low-frequency illumination field was estimated for each slice by Gaussian blurring and used to normalize image intensities, followed by local contrast enhancement. The processed data were then segmented using tUbeNet^23^, a deep-learning framework for 3D vessel segmentation, using the pretrained model. The resulting tUbeNet probability maps (with isotropic voxels) were post-processed by 3D Gaussian filtering (sigma = 0.8). Next, data was normalized to range from 0 to 1 and thresholded using 0.15 cutoff to exclude noise. Small holes inside vessels were filled using connectivity-based morphological operations in 3D and per slice, and small disconnected objects (containing less than 3000 voxels for Podocalyxin and 8000 voxels for αSMA channel) were removed based on connectivity component analysis. The final processed probability maps were then converted to binary masks, which were resampled along the z-axis using nearest-neighbor interpolation, with the scaling factor determined from the original anisotropic voxel dimensions, to obtain isotropic voxels (0.65 ×0.65 ×0.65 µm^3^). Binary masks were skeletonized to a 1-voxel centerline skeleton using the bwskel function (MATLAB 2023b). The skeletonized data sets were converted to mathematical graph format by using custom MATLAB pipeline (See Supplementary Material), and geometrical properties of vessel segments: diameter, length and tortuosity were calculated across arteries, capillaries and veins as follows: 1) Diameter was estimated from the binary vessel mask using the Euclidean distance transform, where the local radius at each centerline voxel was taken as the distance to the nearest background (zero) voxel and converted to diameter (2×radius). Diameter of a segment was computed as mean value of the centerline diameters along that segment. 2) Length was computed as the centerline arc length between two nodes that were either branching or ending points. 3) Tortuosity was computed as the ratio of the length and Euclidian distance between segments terminal nodes. Arterial properties are obtained from binary masks from 568 nm channel staining with αSMA. Capillaries and veins binary masks were obtained from 647 nm channel, stained with Podocalyxin. To avoid double counting, the arterial mask was overlaid on the veins/capillaries mask, and the intersection was removed from the capillary/vein compartment mask. Staining of pial vessels was incomplete, which led to multiple gaps and hollow tubes, thus they were removed from the final analysis for both channels. Capillaries and veins were separated using a diameter threshold of 10 µm (≤10 µm: capillaries, >10 µm: veins). Vascular density (volume fraction) as a function of cortical depth was computed for each sample from selected 3D with homogenous cortical curvature. In samples with a curved pial surface, multiple regions were analyzed and averaged to obtain the final density profile. Density was quantified in consecutive 25 µm bins along the z-direction. Cortical depth was defined in the xy plane as the local trajectory from the pial surface to the GM/WM boundary and was normalized from 0 to 1, where 0 corresponded to the pial surface and 1 to the gray matter/white matter boundary.

**Table 1.** Sizes of regions of interest (ROIs) selected for high resolution imaging.

| Sample name | Width ($\mu\text{m}$) | Hight ($\mu\text{m}$) | Depth ($\mu\text{m}$) |
| --- | --- | --- | --- |
| S1 | 1389 | 3216 | 599 |
| S2 | 3750 | 1964 | 593 |
| S3 | 2030 | 3010 | 458 |
| S4 | 3640 | 1373 | 400 |
| S5 | 2941 | 3732 | 1217 |
| S6 | 2538 | 2532 | 2012 |

## Acknowledgements

We thank Thomas E. Olausson and Nico van den Berg for valuable assistance with image segmentation and data processing, and Serge Dumoulin for constructive discussions.

## Author Contribution

Vanja Ćurčić, Youri Adolfs, and Natalia Petridou designed the study. Vanja Ćurčić and Youri Adolfs designed and performed the experiments and acquired the data. Vanja Ćurčić developed the analysis pipeline and drafted the manuscript. R. Jeroen Pasterkamp provided recources to conduct experiments. Natalia Petridou, Youri Adolfs, Matthias J. P. van Osch, and R. Jeroen Pasterkamp contributed to manuscript revision.

## Materials &Correspondence

All data will be shared upon request. Code will be uploaded to GitHub and shared.

