## Supplementary material for "High-Resolution 3D Mapping of Human Cortical Vasculature"

**5 Supplementary materials**

**5.1 Graph-based representation of vascular data**

The vascular data were converted into a mathematical graph, in which nodes represented vessel bifurcations and endpoints, and edges represented vessel segments connecting pairs of nodes. This representation reduced the vascular network to a lower-dimensional and memory-efficient form while preserving the spatial coordinates and associated diameters of individual vessels. The graph was generated in MATLAB from the binary vessel mask and its corresponding 1-voxel skeleton. After skeletonization and pruning of spurious short branches, branchpoints and endpoints were identified using bwmorph3. Branchpoint clusters were contracted to single representative voxels, and node coordinates were defined as the centroids of endpoint or contracted branchpoint clusters. Node voxels were then removed from the skeleton such that the remaining connected components corresponded to individual vessel segments. Small, connected components were discarded using bwareaopen with a threshold of 5 voxels. For each segment, voxel coordinates, start and end positions, nearest nodes, Euclidean end-to-end distance, segment length, tortuosity, and diameter were computed.

**Supplementary Figure 1**. Optimizing bleaching prior to clearing and comparison of h-iDISCO+ and h-iDISCO+ antigenicity preservation in postmortem human brain samples.

**A - F**) Human brain tissue samples after clearing placed on millimeter paper for scale. Samples were bleached in different conditions prior to clearing. Scale bars 500 µm.

**A - C**) Samples were bleached at 4 degrees Celsius for 7 days and increased concentration of H_2_O_2._ **A**) 5% H_2_O_2_ **B**) 10% H_2_O_2_ and **C**) 15% H_2_O_2._

**D - F**) Samples were bleached at room temperature (RT) and under LED light, on shaking platform, with bleaching lasting for 7 days, and increasing concentration of H_2_O_2._ H_2_O_2_ was refreshed daily. **D**) 5% H_2_O_2_, **E**) 10% H_2_O_2_ and **F**) 15% H_2_O_2._ The final sample (F) showed best optical transparency and 15% of H_2_O_2_ at RT combined with LED light was selected for the final bleaching used in h-iDISCO+.

**G - L**) Representative 100 µm z-projections showing the effects of bleaching conditions on antibody labelling and light penetration. All samples were illuminated with the laser sheet coming from the left side. Boxed regions are shown at higher magnification. Compared with the standard condition (5% H_2_O_2,_ 4 degrees Celsius, overnight), the extended oxidative bleaching step (15% H_2_O_2_, room temperature with LED illumination and on shaking platform, 7 days) improved optical transparency and resulted in more homogeneous light penetration across the tissue without loss of antigenicity.

**G), H**) Podocalyxin staining (concentration 1:200) remained robust after extended bleaching, while signal distribution across the sample became more uniform due to more uniform light penetration.

**I), J**) A similar effect was observed for αSMA staining (concentration 1:200), with preserved labeling and significantly improved light penetration, facilitating imaging of with one-sided illumination.

**K), L**) CD31 staining (concentration 1:65) also showed improved signal distribution after extended bleaching, although the difference was less pronounced due to low specificity and high noise. Due to low signal quality, CD31 was not used in the final protocol.

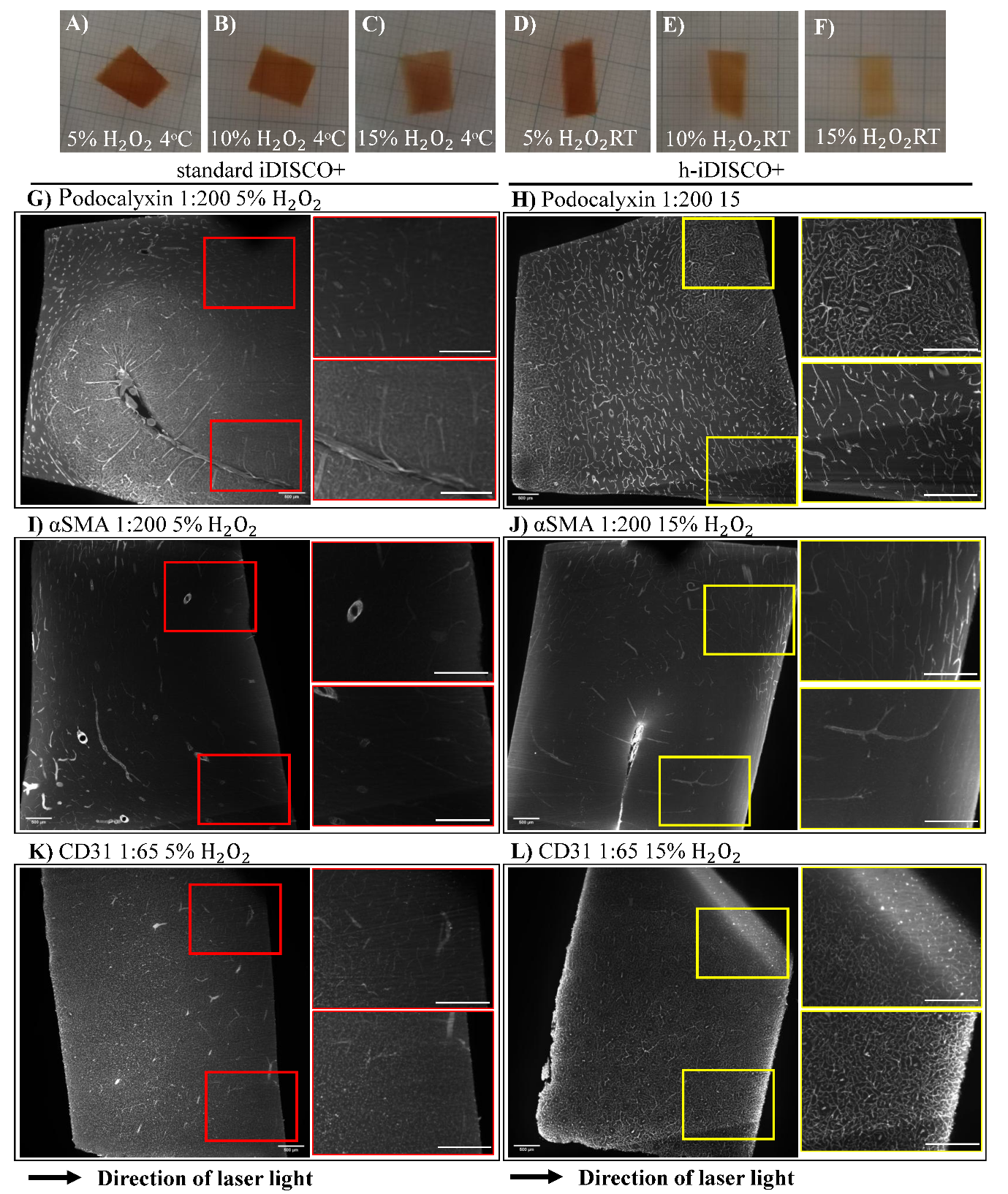

**Supplementary Figure 2**. Optimization of antibody concentrations for vascular labelling in cleared human brain samples.

**A),B**) Podocalyxin staining in whole-mount samples at two antibody concentrations. **A**) At concentration 1:40, signal was strong at the sample border, showing limited antibody penetration and formation or rim around the sample; zoomed insets show antibody accumulation, particularly near the borders (insets 1,3). **B**) At 1:200, Podocalyxin staining showed improved penetration and more uniform vascular labelling throughout the sample.

**C - I**) Assessment of αSMA staining. **C**) Whole-mount staining with αSMA at concentration 1:200 labelled a subset of vessels, with stronger signal in white matter, and fewer labelled branches in grey matter. Additional experiments were performed on 50-µm sections at the same concentration. **D**) αSMA channel. **E**) Overlay of αSMA with the 488-nm autofluorescence channel. **G-I**) Higher magnification views of the boxed regions in D) and E). **G**) Bright circular structures in the αSMA channel correspond to penetrating arteries oriented perpendicular to the section plane. **H**) The same circular structures have low signal in 488 nm channel indicating successful staining. **F**) Lowering the αSMA concentration to 1:1000 still revealed some arteries, but staining quality was reduced compared with the selected 1:200 condition.

**J - P**) Assessment of CD31 staining. **J**) Whole-mount staining with CD31 at concentration 1:65 showed diffuse, non-specific signal. Additional experiments were performed on 50-µm sections at the same concentration. **K**) CD31 channel. **L**) Overlay of CD31 with the 488-nm autofluorescence channel.

**N - P**) Higher magnification views of the boxed regions in K) and L). **N**) CD31 showed non-specific signal, staining both vessels and background, whereas **O**) the autofluorescence channel showed only few vascular structures. **P**) The overlay of CD31 and autofluorescence channel showed that CD31 labelled non-vascular structures, resulting in poor specificity for vascular staining. **M**) Reducing the CD31 concentration to 1:300 resulted in weak labelling, with only a few vessels remaining visible.

These experiments identified Podocalyxin 1:200 and αSMA 1:200 concentrations as the optimal conditions for subsequent experiments, whereas CD31 was excluded because of poor specificity and weak signal.

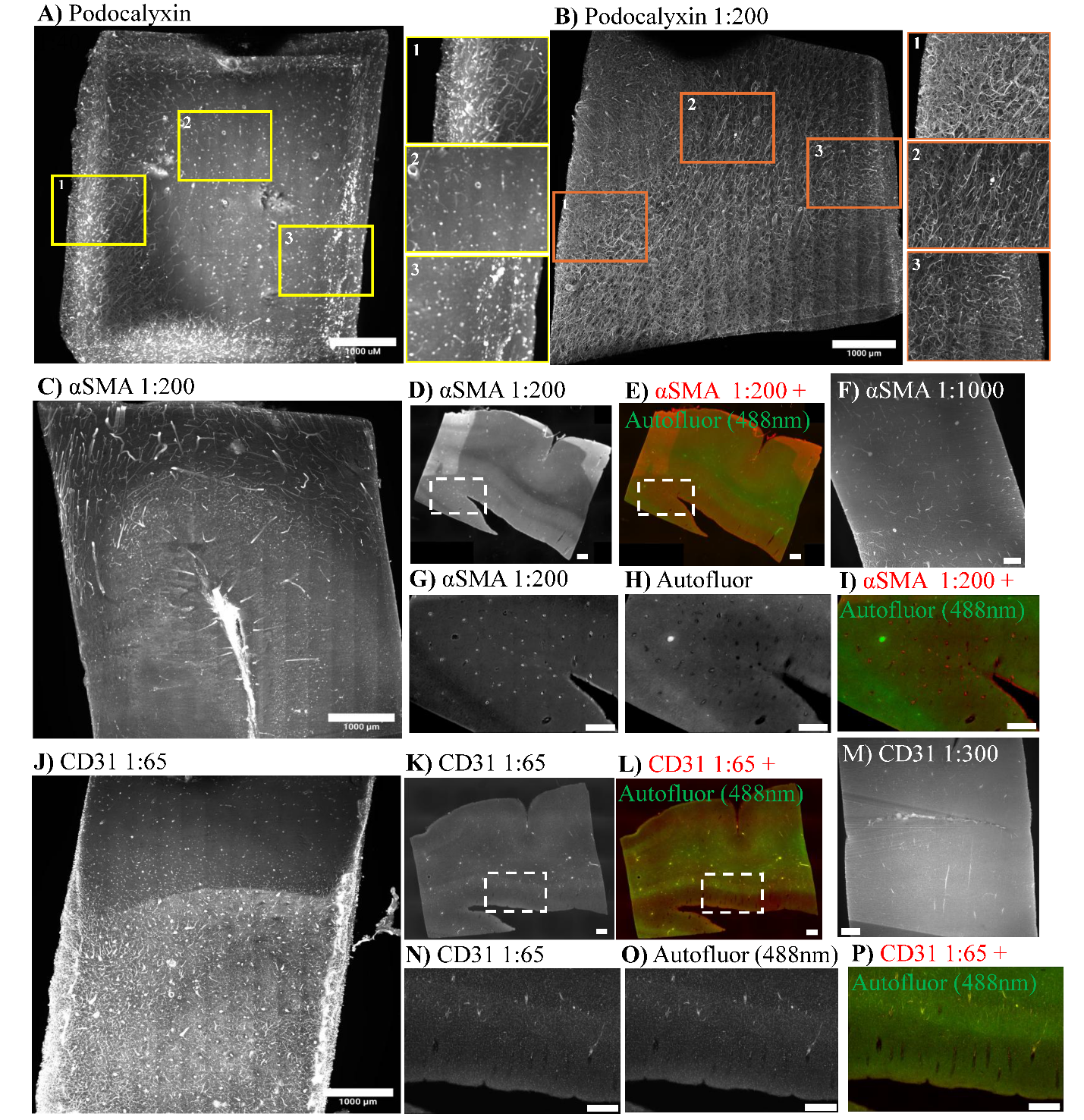
